# Structural modification of naturally-occurring phenolics as a strategy for developing cytotoxic molecules towards cancer cells

**DOI:** 10.1101/2023.03.26.534257

**Authors:** Pedro Olim, Renato B. Pereira, Maria José G. Fernandes, Carolina M. Natal, José R. A. Coelho, A. Gil Fortes, M. Sameiro, T. Gonçalves, David M. Pereira

## Abstract

Natural products belonging to different chemical classes have been established as a promising source of novel anticancer drugs. Several low molecular weight compounds from the classes of monoterpenes, phenylpropanoids and flavonoids were shown to possess anticancer activities in previous studies. In this work, over 20 semisynthetic derivatives of molecules belonging to these classes, namely thymol, eugenol and 6-hydroxyflavanone were synthesized and tested for their cytotoxicity against two human cancer cell lines, namely gastric adenocarcinoma (AGS cells) and human lung carcinoma (A549 cells). An initial screening based on viability assessment was performed in order to identify the most cytotoxic compounds at 100 μM. The results evidenced that two 6-hydroxyflavanone derivatives were the most cytotoxic among the compounds tested, being selected for further studies. Noteworthy, in a general way some of the derivatives synthesized displayed enhanced toxicity when compared with their natural counterparts. Moreover, LDH assay showed that the loss of cell viability was not accompanied by a loss of membrane integrity, thus ruling out a necrotic process. Morphological studies with AGS cells demonstrated chromatin condensation compatible with apoptosis, confirmed by the activation of caspase 3/7. Furthermore, a viability assay on non-cancer human embryonic lung fibroblast cell line (MRC-5) confirmed these two derivatives possess selective anticancer activity.

## 1. Introduction

Thymol and eugenol are the main constituents of the essential oils of thyme (*Thymus vulgaris*) and clove (*Syzygium aromaticum*), respectively. 6-Hydroxyflavanone and other flavanones are widely distributed in plant products, being also the most abundant flavonoid compounds in the peel of citrus fruits.

Plants, like other organisms, have always evolved chemical and biosynthetic tools to further gain an advantage over their natural competitors [1–3]. For this reason, the chemical/metabolic pathways and corresponding products of these organisms have also been subjected to evolutionary pressures. Natural molecules are, for this reason, privileged for their chemical and structural diversity, and for their potential to bind and interact with multiple biological structures and targets, which vastly increases the probability of these molecules exhibiting pharmacological activity [4].

The search for new anticancer drugs has often relied on the diversity and pharmacological potential of natural or semisynthetic small molecules. From the 1940s until 2014, up to 49% of all anticancer drugs approved by the Food and Drug Administration (FDA) were natural products or derived from them [5].

The three parent molecules used in this work are natural products, namely thymol, eugenol, and 6-hydroxyflavanone. The rational for choosing these molecules are twofold: for one, they all belong to different natural product chemical families, namely monoterpenes, phenylpropanoids, and flavonoids, respectively. Second, all these parent molecules have shown some degree of cytotoxic and anticancer activity in previous studies [6–8]. Chemical derivatization has been proven to enhance their cytotoxic activities. A study conducted with β-amino and β-alkoxy alcohols derivatives of eugenol showed these molecules possessed an enhanced but also selective anticancer activity [9]. Similar claims have been made for derivatives of thymol and flavanones [10,11].

In this work, twenty three synthetic derivatives of these three parent natural products were synthesized and tested for their cytotoxic activity. The two best-performing molecules that showed improved cytotoxicity were selected and chosen for further studies. We ultimately aimed to use chemical modification and medicinal chemistry approaches towards the improvement of naturally-occurring phenolic molecules as a strategy for developing cytotoxic anticancer drugs.

## 2. Experimental

### 2.1. Chemicals and standards

All solvents and reagents used in the synthetic work were purchased from Sigma-Aldrich (St. Louis, MO, USA), Fisher Scientific (Geel, Belgium) and PanReac Applichem (Barcelona, Spain. The deuterated solvents were purchased from Eurisotop (Cambridge, England). The TLC analyses were performed on 0.25 mm thick precoated silica plates (Merck Fertigplatten Kieselgel 60F254, Germany), and visualization of the spots was done under UV light. Column chromatography on silica gel was carried out on Merck Kieselgel (230-240 mesh).

Triton X-100, Trypan blue, Dulbecco’s modified eagle medium (DMEM), minimal essential medium (MEM), fetal bovine serum (FBS), 0.25% Trypsin-EDTA, Hanks’ balanced salt solution (HBSS) and penicillin/streptomycin solution (10000 units/mL; 10000 μg/mL, respectively) were purchased from GIBCO (Invitrogen, NY, USA). CytoTox 96® non-radioactive cytotoxicity and caspase-glo® 3/7 assay kits were purchased from Promega Corporation (WI, USA). Staurosporine was obtained from Santa Cruz Biotechnology (TX, USA). QubitTM dsDNA HS Assay Kit for DNA quantification was purchased from Thermo Fisher Scientific (Waltham, MA, USA). Phalloidin-tetramethylrhodamine B isothiocyanate, 4’,6-diamidino-2-phenylindole (DAPI) and dimethyl sulfoxide (DMSO) were from Sigma-Aldrich (Madrid, Spain).

### 2.2. Analytical instruments

The NMR spectra were obtained on a Bruker Avance III at an operating frequency of 400 MHz for ^1^H NMR and 100 MHz for ^13^C NMR using the solvent peak as internal reference at 25 °C. All chemical shifts are given in ppm using *δ* Me4Si = 0 ppm as reference and *J* values are given in Hz. Assignments were made by comparison of chemical shifts, peak multiplicities and *J* values and were supported by spin decoupling-double resonance and bidimensional heteronuclear correlation techniques. The IR spectra were obtained on a Spectrum Two FTIR spectrophotometer PerkinElmer (USA). Mass spectrometry analyses were performed at the “C.A.C.T.I. - Unidad de Espectrometria de Masas”, at University of Vigo, Spain. All melting points were measured on a Stuart Analogue Melting Point Apparatus SMP11 (Cheshire, England).

### 2.3. Synthesis

Compounds **EU1**, **EU2**, **EU4**, **EU5** were synthesised by O-alkylation of eugenol, using 1-bromopropane, 3-bromopropan-1-ol, 1-bromo-3-chloropropane and ethyl 4-bromobutanoate, respectively, with cesium carbonate, by heating at 65°C, in acetonitrile. Compound **EU3** was obtained by reaction of compound **EU2** with acetic anhydride by heating at 65°C. In addition, compound **EU6** resulted from the hydrolysis in aqueous 1M sodium hydroxide/1,4-dioxane, at room temperature of **EU5** [12].

The synthesis of oxiranes **EU7-11** was performed by the reaction of compounds **EU1-5**, respectively, with *m*-chloroperbenzoic acid in dichloromethane, at room temperature [12] Eugenol esters **EU12-16** were synthesized by reaction of eugenol **1** with the corresponding carboxylic acids in dichloromethane, at room temperature, in the presence of 4-dimethylaminopyridine (DMAP) and N,N’-dicyclohexylcarbodiimide (DCC). The synthesis of compounds **EU12** and **EU13** is unpublished in the literature; compounds **EU14** [13,14], and **EU16** [15] have already been obtained, but by different synthetic routes; synthesis of compound **EU15** was previously published [14]. As a result, detail experimental procedure and full characterisation of compounds **EU12-14** and **EU16** are described below.

General procedure for the synthesis of **EU12-14** and **EU16**: a mixture of 4-allyl-2-methoxyphenol (trivialy designated as eugenol), 4-dimethylaminopyridine (DMAP) (0.2-0.62 eq.) and *N,N*-dicyclohexylcarbodiimide (DCC) (1.5 eq.) was added to the corresponding carboxylic acid (1.5 eq.) in dichloromethane (7.5 mL). The reaction mixture was stirred at room temperature for 24 h and monitored by TLC (silica: dichloromethane). At the end of this period, the white suspension obtained was filtered and the liquid phases were washed successively with 1 M hydrochloric acid (2×20 mL), saturated sodium hydrogen carbonate solution (2×20 mL) and water (2×20mL). Finally, after drying with anhydrous sodium sulfate, the organic phases were evaporated under reduced pressure to give the respective ester.

4-Allyl-2-methoxyphenyl quinoline-2-carboxylate **EU12**: starting from 4-allyl-2-methoxyphenol (0.500 g, 3.05 mmol) and using DMAP (0.075 g, 0.61 mmol), DCC (0.944 g, 4.56 mmol) and quinaldic acid (0.793 g, 4.58 mmol), compound **EU12** was obtained as a pink solid (0.224 g; 23%). R*f* = 0.64 (silica: dichloromethane), m.p. = 77–79 °C. FTIR (solid): *v*_max_ 2928, 1739 (C=O), 1638, 1506, 1310, 1292, 1266, 1241, 1200, 1187, 1147, 1125, 1084, 1030, 923, 842, 771 cm^-1^. ^1^H NMR (CDCl_3_, 400 MHz): *δ*_H_ 3.44 (2H, d, *J* = 6.8 Hz, C*H*_2_Ph), 3.83 (3H, s, OC*H*_3_), 5.11-5.17 (2H, m, CH=C*H*_2_), 5.98-6.05 (1H, m, CH=CH_2_), 6.84 (1H, d, *J* = 2.0 Hz, H-3), 6.86 (1H, dd, *J* = 8.0 and 1.2 Hz, H-5), 7.18 (1H, d, *J* = 8.0 Hz, H-6), 7.69 (1H, dt, *J* = 8.0 and 1.2 Hz, H-6 quinoline), 7.82 (1H, dt, *J* = 8.0 and 1.2 Hz, H-7 quinoline), 7.93 (1H, dd, *J* = 8.4 and 1.2 Hz, H-8 quinoline), 8.33-8.38 (2H, m, H-3 and H-5 quinoline), 8.40 (1H, d, *J* = 8.4 Hz, H-4 quinoline) ppm. ^13^C NMR (CDCl_3_, 100 MHz): *δ*_C_ 40.15 (*C*H_2_Ph), 55.87 (OCH_3_), 112.81 (C-3), 116.21 (CH=*C*H_2_), 120.71 (C-5), 122.60 (C-6), 127.56 (C-3 quinoline), 128.80 (C-5 quinoline), 129.52 (C-6 quinoline), 130.30 (C-7 quinoline), 130.69 (C-4a), 131.00 (C-4 quinoline), 137.08 (C-8 quinoline), 137.31 (C-1), 138.35 (*C*H=CH_2_), 139.30 (C-4), 147.43 (C8a), 147.78 (C-2 quinoline), 150.95 (C-2), 162.10 (C=O) ppm. HRMS (ESI) calcd for C_20_H_18_NO_3_ [M+H]^+^ 320.1287, found 320.1290.

4-Allyl-2-methoxyphenyl pyrazine-2-carboxylate **EU13**: starting from 4-allyl-2-methoxyphenol (0.500 g, 3.05 mmol) and using DMAP (0.235 g, 1.92 mmol), DCC (0.944 g, 4.56 mmol) and pyrazinoic acid (0.568 g, 4.58 mmol), compound **EU13** was obtained as a dark yellow solid (0.136 g; 16%). R*f* = 0.51 (silica: dichloromethane), m.p. = 89.0-91.0 °C. FTIR (solid): *v*_max_ 3278, 3073, 2931, 2854, 1757 (C=O), 1651, 1604, 1507, 1301, 1281, 1271, 1196, 1173, 1121, 1107, 1028, 1017, 917, 841, 818, 766 cm^-1^. ^1^H NMR (CDCl_3_, 400 MHz): *δ*_H_ 3.43 (2H, d, *J* = 6.8 Hz, C*H*_2_Ph), 3.82 (3H, s, OCH_3_), 5.11-5.17 (2H, m, CH=C*H*_2_), 5.96-6.03 (1H, m, *C*H=CH_2_), 6.63-6.86 (2H, m, H-3 and H-5), 7.13 (1H, d, *J* = 8.0 Hz, H-6), 8.81 (1H, d, *J* = 1.2 Hz, H-6 pyrazine), 8.82 (1H, d, *J* = 1.2 Hz, H-5 pyrazine), 8.81 (1H, d, *J* = 2.8 Hz, H-3 pyrazine) ppm. ^13^C NMR (CDCl_3_, 100 MHz): *δ*_C_ 40.11 (*C*H_2_Ph), 55.83 (OCH_3_), 112.82 (C-3), 116.30 (*C*H=CH_2_), 120.74 (C-5), 122.30 (C-6), 136.90 (*C*H=CH_2_), 137.65 (C-1), 139.67 (C-4), 143.03 (C-2 pyrazine), 144.60 (C-3 pyrazine), 146.83 (C-5 pyrazine), 147.94 (C-6 pyrazine), 150.69 (C-2), 162.10 (C=O) ppm. HRMS (ESI) calcd for C_15_H_15_N2O_3_ [M+H]^+^ 271.1084, found 271.1082.

4-Allyl-2-methoxyphenyl 4-methoxybenzoate **EU14**: starting from 4-allyl-2-methoxyphenol (0.500 g, 3.05 mmol) and using DMAP (0.075 g, 0.61 mmol), DCC (0.944 g, 4.56 mmol) and anisic acid (0.703 g, 4.62 mmol), compound **EU14** was obtained as a white solid (0.324 g; 36%). R*f* = 0.47 (silica: dichloromethane), m.p. = 88.0-90.0 °C. FTIR (solid): *v*_max_ 2939, 1727 (C=O), 1605, 1578, 1513, 1421, 1283, 1263, 1199, 1183, 1165, 1146, 1118, 1068, 1028, 923, 842, 765 cm^-1^. ^1^H NMR (CDCl_3_, 400 MHz): *δ*_H_ 3.41 (2H, d, *J* = 6.8 Hz, C*H*_2_Ph), 3.81 (3H, s, OCH_3_), 3.90 (3H, s, OC*H*_3_-Ph), 5.10-5.16 (2H, m, CH=C*H*_2_), 5.95–6.03 (1H, m, *C*H=CH_2_), 6.81 (1H, d, *J* = 2.0 Hz, H-3), 6.83 (1H, dd, *J* = 8.0 and 2.0 Hz, H-5), 6.99 (2H, d, *J* = 8.0 Hz, H-3 and H-5 OCH_3_-Ph), 7.06 (1H, d, *J* = 8.0 Hz, H-6), 8.18 (2H, d, *J* = 8.0 Hz, H-2 and H-6 Ph-O*C*H_3_) ppm. ^13^C NMR (CDCl_3_, 100 MHz): *δ*_C_ 40.11 (*C*H_2_Ph), 55.48 (OCH_3_), 55.89 (Ph-OCH_3_), 112.85 (C-5), 113.74 (C-3 and C-5 Ph-OCH_3_), 116.08 (CH=CH_2_), 120.71 (C-3), 121.86 (C-1 Ph-OCH_3_), 122.74 (C-6), 132.39 (C-2 and C-6 Ph-OCH_3_), 137.15 (CH=CH_2_), 138.31 (C-1), 138.86 (C-4), 151.19 (C-2), 163.75 (C-4 Ph-OCH_3_), 164.62 (C=O) ppm. HRMS (ESI) calcd for C_18_H_19_O_4_ [M+H]^+^ 299.1284, found 299.1288.

4-Allyl-2-methoxyphenyl 2-chlorobenzoate **EU16**: starting from 4-allyl-2-methoxyphenol (0.500 g, 3.05 mmol) and using DMAP (0.235 g, 1.92 mmol), DCC (0.944 g, 4.56 mmol) and 2-chlorobenzoic acid (0.716 g, 4.58 mmol), compound **EU16** was obtained as a a yellow oil (0.137 g; 15%). R*f* = 0.73 (silica: dichloromethane). ^1^H NMR (CDCl_3_, 400 MHz): *δ*_H_ 3.41 (2H, d, *J* = 6.8 Hz,*C*H_2_Ph), 3.81 (3H, s, OCH_3_), 3.90 (3H, s, OC*H*_3_-Ph), 5.10–5.16 (2H, m, CH=*C*H_2_), 5.95–6.03 (1H, m, *C*H=CH_2_), 6.81 (1H, d, *J* = 2.0 Hz, H-3), 6.83 (1H, dd, *J* = 8.0 and 2.0 Hz, H-5), 6.99 (2H, d, *J* = 8.0 Hz, H-3 and H-5 OCH_3_-Ph), 7.06 (1H, d, *J* = 8.0 Hz, H-6), 8.18 (2H, d, *J* = 8.0 Hz, H-2 and H-6 Ph-OCH_3_) ppm. ^13^C NMR (CDCl_3_, 100 MHz): *δ*_C_ 40.11 (*C*H_2_Ph), 55.48 (OCH_3_), 55.89 (Ph-OCH_3_), 112.85 (C-5), 113.74 (C-3 and C-5 Ph-OCH_3_), 116.08 (*C*H=CH_2_), 120.71 (C-3), 121.86 (C-1 Ph-OCH_3_), 122.74 (C-6), 132.39 (C-2 and C-6 Ph-OCH_3_), 137.15 (CH=CH_2_), 138.31 (C-1), 138.86 (C-4), 151.19 (C-2), 163.75 (C-4 Ph-OCH_3_), 164.62 (C=O) ppm. HRMS (ESI) calcd for C_18_H_19_O_4_ [M+H]^+^ 299.1284, found 299.1288.

Compounds **TH1-3** [16] and **TH4,5** were synthesised by O-alkylation of tymol (trivial name for 2-isopropyl-5-methylphenol), using 3-bromopropan-1-ol, 1-bromo-3-chloropropane, 1-bromopropane, 1-bromododecane and 1-bromoctane,_respectively, with cesium carbonate, by heating at 65°C, in acetonitrile. The synthesis of compounds **TH4,5** is unpublished, and therefore detail experimental procedure and full characterisation are described below.

General procedure for the synthesis of compounds **TH4,5**: to a solution of 2-isopropyl-5-methylphenol **2** (1 equiv) in acetonitrile (4 mL), the corresponding alkyl halide (1.1 equiv) and cesium carbonate (5 equiv) were added, and the resulting mixture was heated at 65 °C for 5 h. The progress of the reaction was monitored by TLC (light petroleum). The excess of base was filtered and the solvent was evaporated to give the corresponding compound.

2-(Dodecyloxy)-1-isopropyl-4-methylbenzene **TH4**: starting from 2-isopropyl-5-methylphenol (0.100 g, 0.67mmol) and using 1-bromododecane (0.176 mL, 0.73 mmol), compound **TH4** was obtained as a light yellow oil (0.197g, 93% yield). R*f* = 0.82 (silica: light petroleum). ^1^H NMR (CDCl_3_, 400 MHz): *δ*_H_ 1.05 (3H, t, *J* = 6.8 Hz, OCH_2_CH_2_CH_2_(CH_2_)_8_C*H*_3_), 1.37 (6H, d, *J* = 6.0 Hz, CH(CH_3_)_2_), 1.41-1.56 (16H, m, OCH_2_CH_2_CH_2_(CH_2_)_8_CH_3_), 1.61-1.68 (2H, m, OCH_2_CH_2_CH_2_(CH_2_)_8_CH_3_), 1.92-1.99 (2H, m, OCH_2_CH_2_CH_2_(CH_2_)_8_CH_3_), 2.50 (3H, s, PhC*H*_3_), 3.41-3.51 (1H, m, *C*H(CH_3_)_2_), 4.09 (2H, t, *J* = 6.4 Hz, OC*H*_2_CH_2_CH_2_(CH_2_)_8_CH_3_), 6.80 (1H, d, *J* = 1.6 Hz, H-3), 6.87 (1H, dd, J = 7.6 Hz, *J* = 1.6 Hz, H-5), 7.23 (1H, d, *J* = 7.6 Hz, H-6) ppm. ^13^C NMR (CDCl_3_, 100 MHz): *δ*_C_ 14.11 (O(CH_2_)_11_*C*H_3_), 21.32 (Ph*C*H_3_), 22.73 (CH_2_), 22.75 (CH(*C*H_3_)_2_), 26.25 (CH_2_), 26.67 (*C*H(CH_3_)_2_), 29.41 (2×CH_2_), 29.49 (CH_2_), 29.66 (2×CH_2_), 29.70 (CH_2_), 29.73(CH_2_), 31.97 (CH_2_), 67.78 (O*C*H_2_(CH_2_)_10_CH_3_), 112.12 (C-3), 120.84 (C-5), 125.76 (C-6), 134.03 (C-4), 136.11 (C-1), 156.23 (C-2) ppm. HRMS (ESI) calcd for C_22_H_39_O [M+H]^+^ 319.3001, found 319.3003.

1-Isopropyl-4-methyl-2-(octyloxy)benzene **TH5**: starting from 2-isopropyl-5-methylphenol (0.120 g, 0.80 mmol) and using 1-bromoctane (0.152 mL, 0.88 mmol), compound **TH5** was obtained as a light yellow oil (0.190 g, 91% yield). R*f* = 0.84 (silica: light petroleum). ^1^H NMR (CDCl_3_, 400 MHz): *δ*_H_ 1.09 (3H, t, *J* = 7.2 Hz, OCH_2_CH_2_CH_2_(CH_2_)_4_C*H*_3_), 1.40 (6H, d, *J* = 6.8 Hz, CH(C*H*_3_)_2_), 1.48-1.59 (8H, m, OCH_2_CH_2_CH_2_(C*H*_2_)_4_CH_3_), 1.67 (2H, quint, *J* = 7.2 Hz, OCH_2_CH_2_C*H*_2_(CH_2_)_4_CH_3_), 1.94-2.01 (2H, m, OCH_2_C*H*_2_CH_2_(CH_2_)_4_CH_3_), 2.49 (3H, s, PhC*H*_3_), 3.44-3.54 (1H, m, CH(C*H*_3_)_2_), 4.11 (2H, t, *J* = 6.4 Hz, OCH_2_CH_2_CH_2_(CH_2_)_4_CH_3_), 6.83 (1H, d, *J* = 1.6 Hz, H-3), 6.90 (1H, dd, *J* = 7.6 e 1.6 Hz, H-5), 7.26 (1H, d, *J* = 7.6 Hz, H-6) ppm. ^13^C NMR (CDCl_3_, 100 MHz): *δ_C_* 14.09 (O(CH_2_)_7_CH_3_), 21.31 (PhC*H*_3_), 22.69 (CH2), 22.73 (CH(C*H*_3_)_2_), 26.24 (CH_2_), 26.66 (CH(C*H*_3_)_2_), 29.30 (CH_2_), 29.35 (CH_2_), 29.48 (CH_2_), 31.86 (CH_2_), 67.76 (OCH_2_(CH_2_)_6_CH_3_), 112.11 (C-3), 120.83 (C-5), 125.75 (C-6), 134.02 (C-1), 136.10 (C-4), 156.22 (C-2) ppm. HRMS (ESI) calcd for C_18_H_31_O [M+H]^+^ 263.2375, found 263.2377.

6-(3-Chloropropoxy)-2-phenylchroman-4-one **HF1** and 2-phenyl-6-propoxychroman-4-one **HF2** were obtained through the alkylation reaction of 6-hydroxyflavanone, trivial name for 6-hydroxy-2-phenylchroman-4-one, in basic medium, at room temperature using cesium hydroxide and tetrabutylammonium iodide (TBAI), in acetonitrile, with 1-bromo-3-chloropropane and 1-bromopropane, respectively. The synthesis of **HF1** and **HF2** are unpublished in the literature, and therefore detailed experimental procedure and characterisation of these compounds are described.

Synthesis of 6-(3-chloropropoxy)-2-phenylchroman-4-one **HF1**: to a solution of 7,6-hydroxy-2-phenylchroman-4-one (0.050 g, 2 mol, 1 equiv) in acetonitrile (3 mL) was added cesium hydroxide (0.066 g, 0.4 mmol, 2 equiv), TBAI (0.082 g, 0.2 mmol, 1.1 equiv) and 1-bromo-3-chloropropane (0.023 mL, 0.2 mmol, 1.1 equiv). The reaction mixture was stirred at room temperature for 24 h, and monitored by TLC in DCM. After this period, the reaction mixture was filtered, the solid was washed with acetonitrile and the solvent was evaporated under reduced pressure to give a residue that was purified by column chromatography (petroleum ether and DCM, mixtures of increasing polarity, as eluent). Compound **HF1** was obtained as a light brown solid (0,012g, 19%), R*f* = 0.85 (silica: DCM), ^1^H NMR (CDCl_3_, 400 MHz): *δ_H_* 2.23-2.28 (2H, m, OCH_2_C*H*_2_CH_2_Cl), 2.86-2.91 (1H, m, H-3), 3.04-3.11 (m, 1H, H-3), 3.75 (2H, t, *J* = 8 Hz, OCH_2_C*H*_2_CH_2_Cl), 4.14 (2H, t, *J* = 8 Hz, OCH_2_C*H*_2_CH_2_Cl), 5.43-5.47 (1H, m, H-2), 7.01 (1H, d, *J* = 8.8 Hz, H-8), 7.13 (1H, dd, *J* = 8.8 and 3.2 Hz, H-7), 7.38-7.50 (6H, m, Ph and H-5) ppm. ^13^C NMR (CDCl_3_, 100 MHz): *δ_C_* 32.15 (OCH_2_C*H*_2_CH_2_Cl), 41.37 (OCH_2_C*H*_2_CH_2_Cl), 44.53 (C-3), 64.93 (OCH_2_C*H*_2_CH_2_Cl), 79.69 (C-2), 108.46 (C-5), 119.45 (C-8), 120.78 (C-4a), 125.58 (C-7), 126.11 (C-2 Ph and C-6 Ph), 128.72 (C-4 Ph), 128.81 (C-5 Ph and C-3 Ph), 138.78 (C-1 Ph), 153.30 (C-6), 156.34 (C-8a), 191.96 (C-4) ppm. HRMS (ESI) calcd for C_18_H_18_ClO_3_ [M+H]^+^ 317.0944, found 317.0946.

Synthesis of 2-phenyl-6-propoxychroman-4-one **HF2**: to a solution of 6-hydroxy-2-phenylchroman-4-one (0.100 g, 0.4 mmol, 1 equiv) in acetonitrile (4 mL) was added cesium hydroxide (0.129 g, 0.9 mmol, 2 equiv), TBAI (0.165 g, 0.4 mmol, 1.1 equiv) and 1-bromopropane (0.045 mL, 0.5 mmol, 1.1 equiv). The reaction mixture was stirred at room temperature for 3 days and monitored by TLC in DCM. After this period, the reaction mixture was filtered, the solid was washed with acetonitrile and the solvent was evaporated under reduced pressure to give a residue that was purified by column chromatography (petroleum ether and DCM, mixtures of increasing polarity, as eluent). Compound **HF2** was obtained as a brown solid (0,018 g; 15%). ^1^H NMR (CDCl_3_, 400 MHz): *δ_H_* 1.05 (3H, t, *J* = 7.6 Hz, OCH_2_CH_2_C*H*_3_), 1.77-1.86 (2H, m, OCH_2_CH_2_C*H*_3_), 2.87-2.92 (1H, m, H-3), 3.04-3.12 (1H, m, H-3), 3.94 (2H, t, *J* = 6.8 Hz, OCH_2_CH_2_C*H*_3_), 5.30-5.47 (1H, m, H-2), 7.00 (1H, d, *J* = 8.8 Hz, H-8), 7.14 (1H, dd, *J* = 9.2 and 3.2 Hz, H-7), 7.36 (1H, d, *J* = 3.2 Hz, H-5), 7.39-7.47 (5H, m, H-Ph) ppm. ^13^C NMR (CDCl_3_, 100 MHz): *δ_C_* 10.47 (OCH_2_CH_2_C*H*_3_), 22.50 (OCH_2_CH_2_C*H*_3_), 44.61 (C-3), 70.17 (OCH_2_CH_2_C*H*_3_), 79.68 (C-2), 108.25 (C-5), 119.32 (C-8), 120.75 (C-4a), 125.78 (C-7), 126.12 (Ph-C2 and Ph-C6), 128.70 (Ph-C4), 128.81 (Ph-C3 and Ph-C5), 138.88 (Ph-C1), 153.75 (C-6), 156.12 (C-8a), 192.11 (C-4) ppm. HRMS (ESI) calcd for C_18_H_19_O_3_ [M+H]^+^ 283.1334, found 283.1332.

All compounds synthesised and used in the present studies had a purity higher than 95% according to the ^1^H NMR spectra.

### 2.4. Stock and working solutions

All compounds (natural and semisynthetic derivatives) were prepared in DMSO at a concentration of 80 mM (stock solution). A set of working solutions were prepared thereafter at 100 μM, by diluting 800 times the previous stock in cell culture medium.

### 2.5. Cell culture

AGS and A549 cells were cultured as monolayers in DMEM, and MRC-5 in MEM, in both cases supplemented with 10% FBS and 1% penicillin/streptomycin at 37°C, in a humidified atmosphere of 5% CO_2_.

For subculture, cells were washed with HBSS, incubated with 0.25% Trypsin-EDTA solution (Sigma-Aldrich) for 3 min at 37°C, resuspended in culture medium and centrifuged at 1300 rpm for 3 min. The supernatant was removed, and the cell pellet was resuspended in 4 mL of culture medium. Cell passages were kept low for all cell lines, with a maximum of 12 passages.

### 2.6. Viability assessment

For the assessment of viability, a tetrazolium dye-based method was used. AGS, A549 and MRC-5 cells were cultured in 96-well plates at a density of 1.5×10^4^, 1.0×10^4^ and 2.0×10^4^ cells *per* well, respectively, allowed to attach for 24 h, and then exposed to the molecules under study for 24 h.

For this method, the tetrazolium dye used was 3-(4,5-dimethylthiazol-2-yl)-2,5-diphenyltetrazolium bromide (MTT), which is converted to (*E,Z*)-5-(4,5-dimethylthiazol-2-yl)-1,3-diphenylformazan (formazan) by mitochondrial reductase in viable cells. After the incubation period, MTT (0.5 mg/mL final concentration) was added to each well and the plate incubated for 2h at 37 °C. Formazan crystals were dissolved with a DMSO:isopropanol mixture (3:1) and then quantified spectrophotometrically at 560 nm in a microplate reader (Multiskan ASCENT, MA, USA). The results correspond to the mean ± standard error of the mean (SEM) of at least three independent experiments performed in triplicate and are expressed as percentage of the untreated control cells.

### 2.7. LDH assay

LDH is a cytosolic enzyme and although it is secreted, its most concentrated in the intracellular environment, thus, a significant increase of LDH in culture medium is a good marker of cell lysis and necrosis. To assess the effect on membrane integrity, cells were cultured as described above for the MTT assay. Then, after 24 h expousure to HF1 and HF2, cell culture media was collected and LDH release was assessed, using a commercial kit (CytoTox96^®^, Promega). The extracellular LDH in the media can be quantified by a coupled enzymatic reaction in which LDH catalyzes the conversion of lactate to pyruvate via NAD^+^ reduction to NADH. Oxidation of NADH by diaphorase leads to the reduction of a tetrazolium salt to a formazan product that can be measured spectrophotometrically at 490 nm, in microplate reader (Multiskan ASCENT). Triton X-100 at 1% was used as positive control for cell lysis. All the results correspond to the fold-increase of absorbance in treated versus untreated cells.

### 2.8. Morphological assessment

For morphological studies, AGS cells were cultured in 96-well plates at the same density used for viability experiments, in the presence of HF1 and HF2 at a concentration of 100 μM. After incubation, cells were washed with HBSS and fixed with 10% formalin solution for 20 min, at room temperature. Phalloidin and DAPI (0.25 μg/mL) were added, and cells were stained for 30 min at room temperature and washed with HBSS, as we have described before [17]. Images were acquired in an inverted Eclipse Ts2R-FL (Nikon) equipped with a Retiga R1 camera and an S Plan Fluor ELWD 20x DIC N1 objective. Images were analyzed with Fiji [18].

### 2.9. DNA quantification

Cells were cultured at the same density described for the MTT assay, in the presence of the molecules under study. After incubation, the culture medium was replaced by 50 μL of ultra-pure water, plates being incubated for 30 min at 37°C and immediately frozen at −80°C. DNA quantification was performed in a triplicate pool using a QubitTM 1X dsDNA HS Assay Kit according to manufacturer’s instructions [19].

### 2.10. Caspase activity

AGS and A549 cells were cultured in 96-well white plates, in the same conditions described above for the MTT assay. After this period, cells were incubated with the test molecules (HF1 and HF2) at 100 μM during 24 h, at 37 °C. Then, 60 μL of cell culture medium was removed from each well, and 40 μL of caspase 3/7 substrate (Promega Corporation, Fitchburg, WI, USA) was added to the wells. Luminescence was determined after incubation for 45 min in a microplate reader (CytationTM 3, BioTek). Results correspond to the fold-increase of luminescence in treated vs untreated cells of three independent experiments, performed in duplicate. Staurosporine (100 nM) was used as positive control for caspase activation and results were normalized to DNA content to account for possible variations in cell mass caused by cell death, as we have described before [19].

### 2.11. Statistical analysis

All data was analyzed using GraphPad Prism 8.4.3 Software (San Diego, CA, USA). Data is expressed as mean ± SEM. Statistical significance between control and positive controls was analyzed by unpaired Student’s t test. Statistical significance between control and cells treated with a set of tested compounds (natural product and derivatives) were analyzed by ANOVA multiple comparisons. For the calculation of IC25 values, a third order and second order polynomial function were used for **HF1** and **HF2**, respectively. Values of p < 0.05 were considered statistically significant.

### 2.12. Visualization

Chemical structures were drawn using rdKit [20], having SMILES as inputs. Figures 2-4 and 6-7 were created using GraphPad Prism 8.4.3 Software (San Diego, CA, USA)

**Figure 1.**
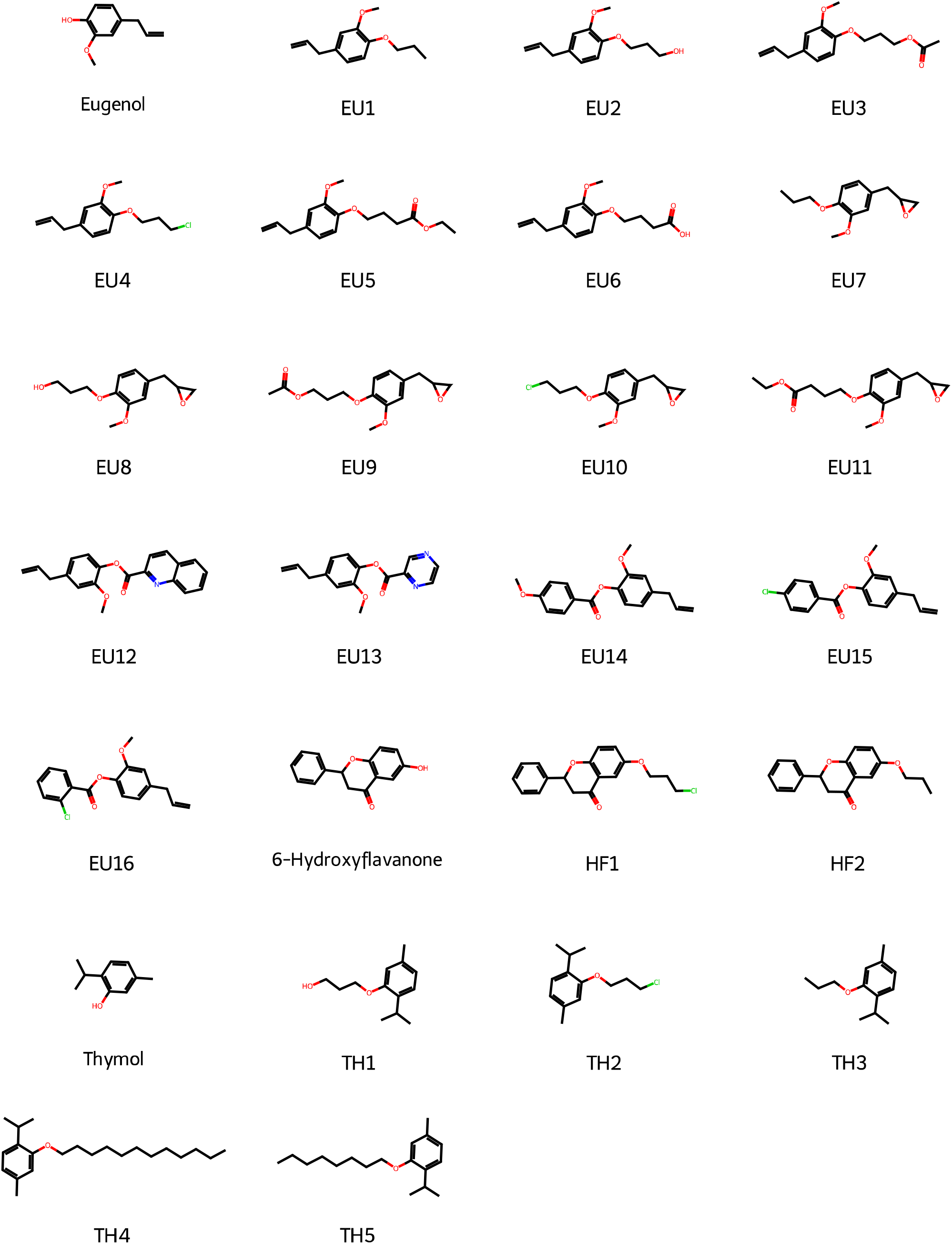
Chemical structures of all the molecules under study. **EU1-16**: eugenol derivatives; **HF1,2**: 6-hydroxyflavanone derivatives; **TH1-5**: thymol derivatives.

**Figure 2.**
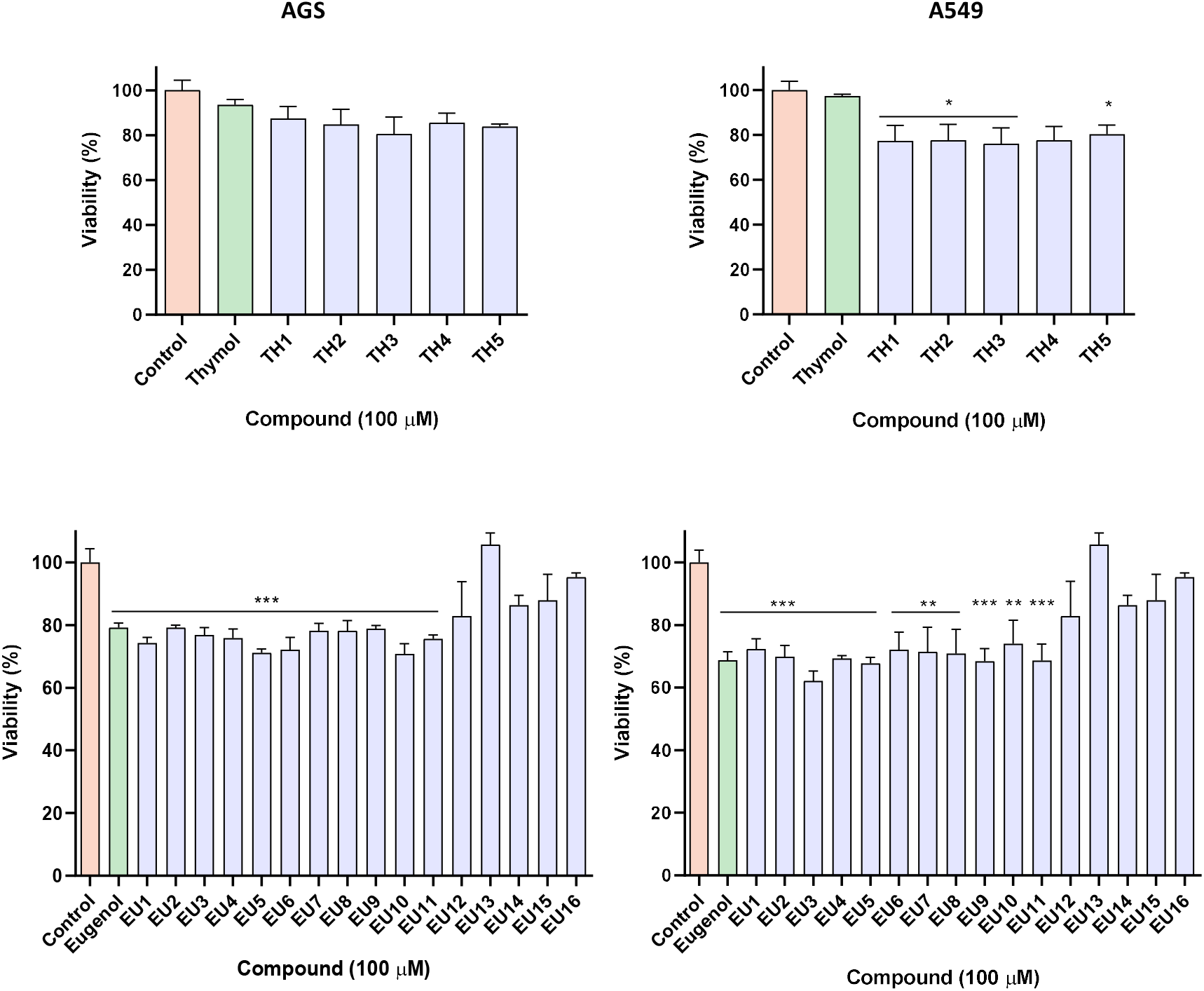
Viability of AGS and A549 cells exposed to thymol (top row) and eugenol (bottom row) and their respective derivatives for 24h based on MTT reduction. Parent molecules are depicted in green while semi-synthetic derivatives are presented in blue. **TH1-5**: thymol derivatives; **EU1-16**: eugenol derivatives. * *p < 0.05, **p< 0.01, ***p < 0.001.*

**Figure 3.**
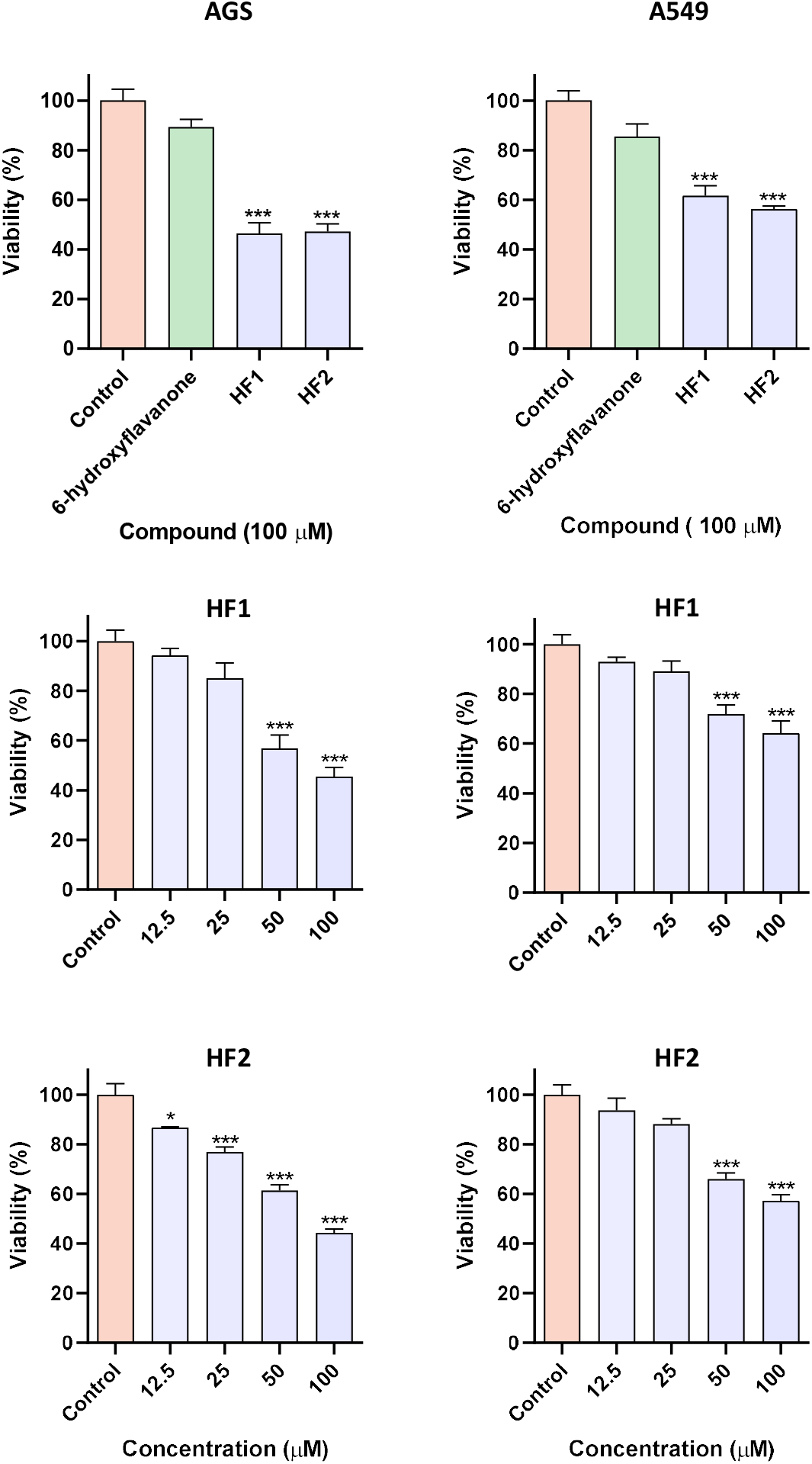
Top row: viability of AGS and A549 cells exposed to 6-hydroxyflavanone and its derivatives (**HF1,2**) for 24h based on MTT reduction. Middle row: viability of AGS and A549 cells exposed to **HF1** in the 12.5-100 μM range. Bottom row: viability of AGS and A549 cells exposed to **HF2** in the 12.5-100 μM range. * *p < 0.05, ***p < 0.001.*

**Figure 4.**
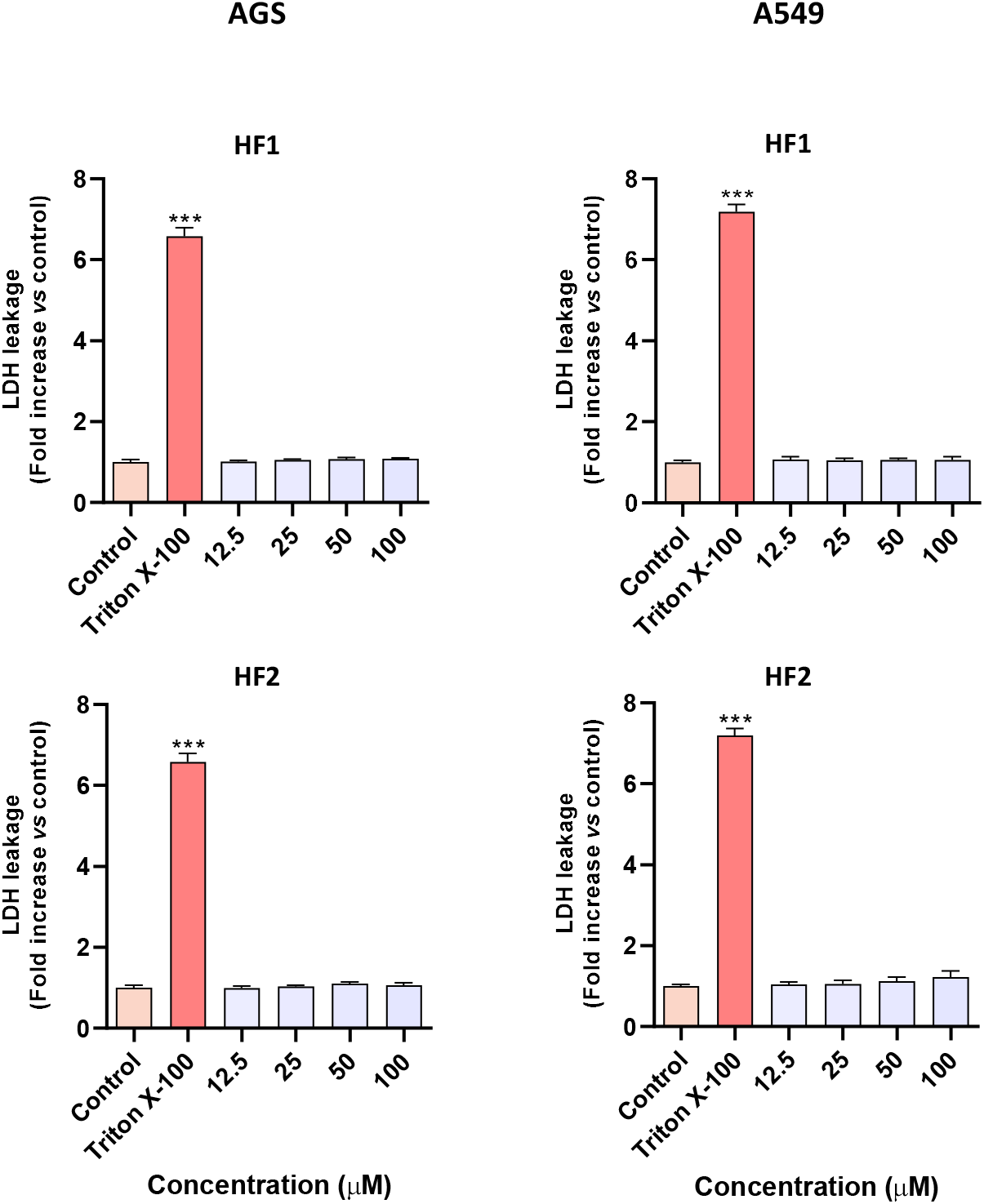
Fold increase in LDH leakage for AGS and A549 cells exposed to **HF1** and **HF2** in the 12.5-100 μM range. Triton X-100 at 1% was used as positive control for membrane lysis. *** *p < 0.001.*

## 3. Results & Discussion

### 3.1. Screening of cytotoxicity

It is clear in the literature that several naturally occurring molecules display a set of pharmacological properties that render them interesting as a starting point for new drugs. Among natural products, phenolics are from the most reported molecules owing to their reactive chemistry that in turn translates frequently into bioactive molecules. Phenolic structures have the potential for strong interations with proteins due to their hydrophobic benzenoid rings and hydroxyl groups which can be involved in hydrogen bonding [21].

In this work, we selected three different phenolic molecules as starting point for the obtention of derivatives for subsequent biological assessment, each of which representing a distinct group of phenolics. The choice had in mind not only previously published data regarding cytotoxic effect of the parent molecules, but was an attempt to select structurally simple molecules in order to facilitate chemical modification and afford cost-effective molecules. In particular, we chose a monoterpene (thymol), phenylpropanoid (eugenol) and flavonoid (6-hydroxyflavanone). These molecules share a number of properties that make them good candidates for semi-synthesis, including inexpensive parent molecule and reactive functional groups.

For each of these parent molecules, several derivatives were prepared, as highlighted in **Figure 1**.

All the molecules synthesized, as well as the parent compounds, were initially evaluated for their impact upon the viability of A549 and AGS cancer cell lines. Considering the high number of molecules, a single concentration of 100 μM was established for testing, which allowed the comparison of the cytotoxic activity of all compounds.

In the case of thymol, the parent molecule was devoid of activity in both cell lines and all derivatives tested displayed toxicity, albeit only in the A549 cell line, however the degree of activity was low (around 20%), reason for which they were deemed devoid of interest for subsequent evaluation (**Figure 2**). In previous studies thymol has shown only moderate cytotoxic activity, a study conducted on AGS cells showed that thymol at 100 μM produced a 10% drop in cell viability after 24h. In other cell lines, such as HT-29 and HCT-116, thymol showed no cytotoxic activity at this concentration [10].

Eugenol was the starting molecule that displayed some degree of toxicity towards cancer cells, namely ca. 20%, with the activity being similar towards both cell lines evaluated (**Figure 2**). As shown in the same figure, while all derivatives obtained were also cytotoxic, they did not exceed the potency of the parent molecule, reason for which they were dropped, and no further studies were conducted. Eugenol, at a concentration of 100 μM, reduced the viability of A549 cells by 20%, which is consistent with other previous studies [11].

The case was different with 6-hydroxyflavanone, which was inactive towards both cell lines while derivatives **HF1** (6-(3-chloropropoxy)flavanone) and **HF2** (6-propoxyflavanone) displayed a marked increase in potency, causing a reduction of cell viability of around 50% (**Figure 3**). For this reason, we selected these molecules for further studies. A similar study on HeLa cells reported a 40% decrease in viability of cells exposed to a concentration of 100 μM of 6-hydroxyflavanone for 48h. The same study showed that 6-propionoxyflavanone, an esterified derivative of 6-hydroxyflavanone, exhibited enhanced cytotoxicity over the parent molecule [22].

### 3.2. The cytotoxic effect of 6-(3-chloropropoxy)-flavanone and 6-propoxyflavanone is dose-dependent and is necrosis-independent

Considering the results found earlier, we were interested in assessing if the effect of 6-(3-chloropropoxy)-flavanone and 6-propoxyflavanone could be concentration-dependent.

To this end, a second viability assessment, similar to that described previously, was performed using several concentrations in the 12.5-100 μM range. The results in **Figure 3** clearly show that both compounds elicit a dose-dependent cytotoxic response in both cancer cell lines. This concentration-dependent behavior has been previously reported for the parent molecule, 6-hydroxyflavanone, and also for 6-propionoxyflavanone, an esterified derivative [22]. In AGS cells, 6-propoxyflavanone was more potent than 6-(3-chloropropoxy)-flavanone at lower doses. In an attempt to calculate the IC50, compounds were also incubated at a concentration of 200 μM, however, this resulted in the insolubility and precipitation in the culture medium, reason for which it was not possible to assess viability at this concentration. For this reason, we chose to calculate the IC25 instead (**HF1**: 34.61 μM and **HF2**: 27.76 μM).

In light of these results, we investigated if these derivatives could be exerting their effect by eliciting necrosis, a process of cell death more commonly linked to pathological processes and characterized by the rapid permeabilization of the cellular membrane, releasing its contents into the extracellular space. Thus, this process of cell death may elicit an inflammatory response, attracting leukocytes and phagocytes [23]. Considering that necrosis-eliciting molecules are frequently devoid of interest for the clinical set, we deemed it important to confirm if this process could be taking place early on in the study. To assess the integrity of the cellular membrane, we searched for the presence of the intracellular enzyme LDH in the culture media, the results being displayed in **Figure 4**. Triton X-100 was used as a positive control for this assay due to its tensioactive properties that cause plasma membrane disruption and leakage of cytoplasmic content, including LDH.

At the concentrations tested, none of the molecules under study caused any evidence of loss of membrane integrity or cell lysis (LDH negative). Considering that the same molecules elicited a marked loss of cell viability at the same concentration, we hypothesized that a process of programmed cell death could be taking place. A study on the cytotoxic and anticancer effects of 6-hydroxyflavanone and C4’, C6 and C7 substituted hydroxylated and methoxylated derivatives on HL-60 cells (peripheral blood lymphocytes) suggested these molecules elicit apoptosis. These derivatives (incubated for 12 h at a concentration of 40 μM) increased the levels of several apoptotic markers such as caspase-3, Bax and cleaved PARP [24]. 6-Hydroxyflavanone and 6-propionoxyflavanone also augmented the anticancer effect of tumor necrosis factor-related apoptosis-inducing ligand (TRAIL), a promising anticancer agent in the TNF superfamily due to its selective effect on cancer cells. As more tumors become resistant to TRAIL-mediated death, drugs capable of sensitizing cells to its effect are much needed. Flavonoids, particularly C6 hydroxylated flavanones, have shown to do so [22].

#### 3.3.6-(3-Chloropropoxy)-flavanone and 6-propoxyflavanone activate effector caspase-3

With necrosis ruled out, we further investigated to putative mechanism behind the cytotoxic effect of 6-(3-chloropropoxy)-flavanone and 6-propoxyflavanone. Among the multiple pathways for organized cell death currently known, apoptosis is the most widely described [25]. Cellular contents, such as DNA, are destroyed in a process initiated by caspases and during which cell membranes retain their structural integrity, thus preventing further immunological response from occurring. As cells are destroyed, they shrink and are split into fragments referred to as apoptotic bodies which can be detected *via* microscopy [26]. These apoptotic bodies, as well as other characteristics such as pyknotic nuclei, can be visualized under a fluorescence microscope when proper stains are used. As shown in **Figure 5**, AGS incubated with either **HF1** or **HF2** showed traits of condensed chromatin in addition to lower cell density, which could be compatible with with a process of organized cell death. In the case of A549 cells, lower cell density was detected, however without appreciable incidence of chromatin condensation (data not shown), which may be compatible with a cytostastic effect.

**Figure 5.**
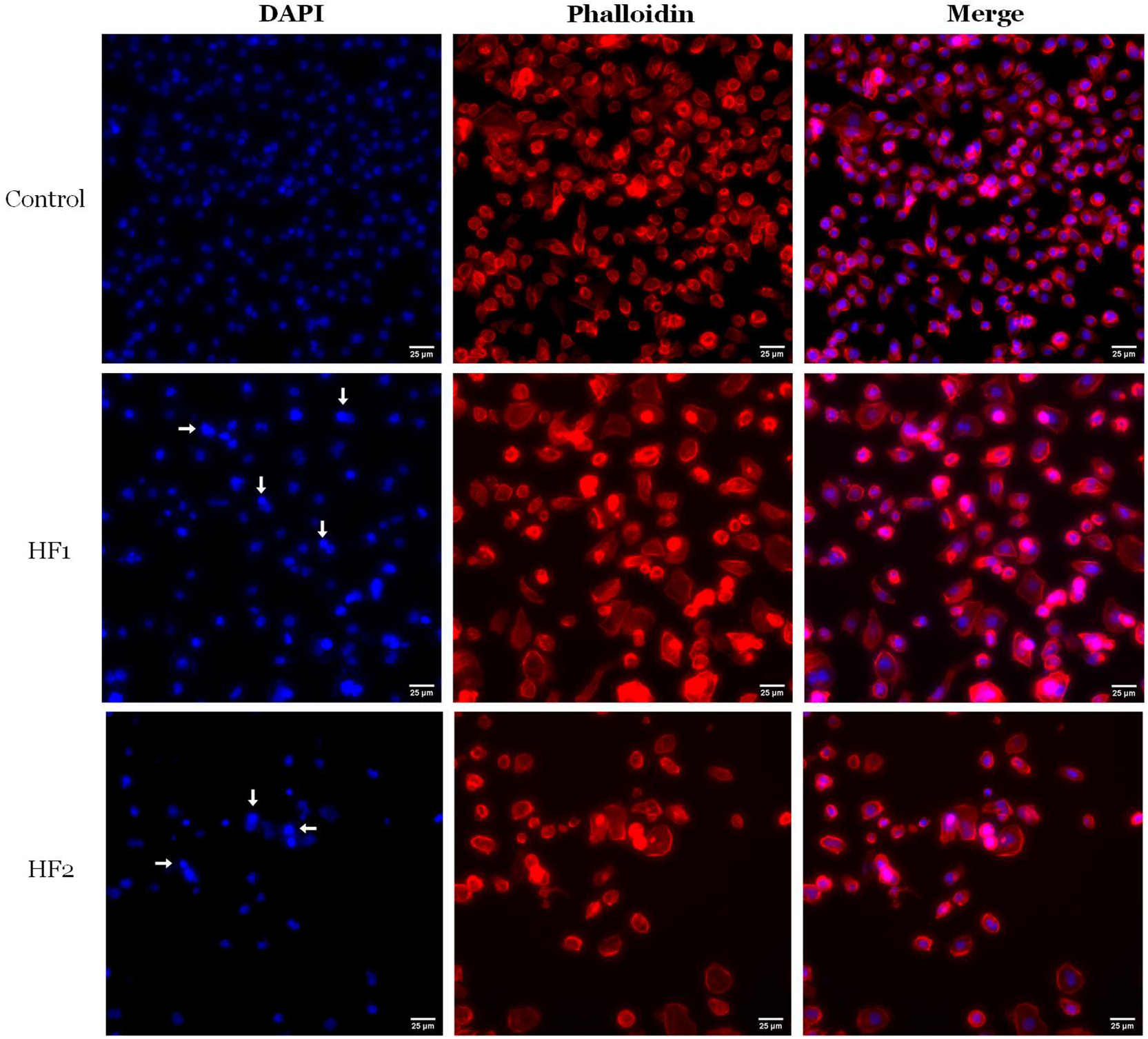
Morphology of AGS cells exposed to compounds **HF1-2** (100 μM) after 24 h of incubation (S Plan Fluor ELWD 20× DIC N1 objective). DAPI (chromatin) and phalloidin (actin). White arrows indicate cells exhibiting condensed chromatin.

To further back up this evidence of apoptosis, another assay was performed, this time to evaluate the activation of effector caspase-3 directly. Considering that caspase activation was studied using a concentration of 6-(3-chloropropoxy)-flavanone and 6-propoxyflavanone that caused marked cytotoxicity, we normalized all results for DNA content in order to rule out the potential interference of cell densities distinct from the control.

As the results presented **Figure 6** show, there was a significant increase in caspase activation, at least for AGS cells, following exposure to 6-(3-chloropropoxy)-flavanone and 6-propoxyflavanone. This further confirms that these compounds induce apoptosis, in agreement from morphological data from **Figure 5**.

**Figure 6.**
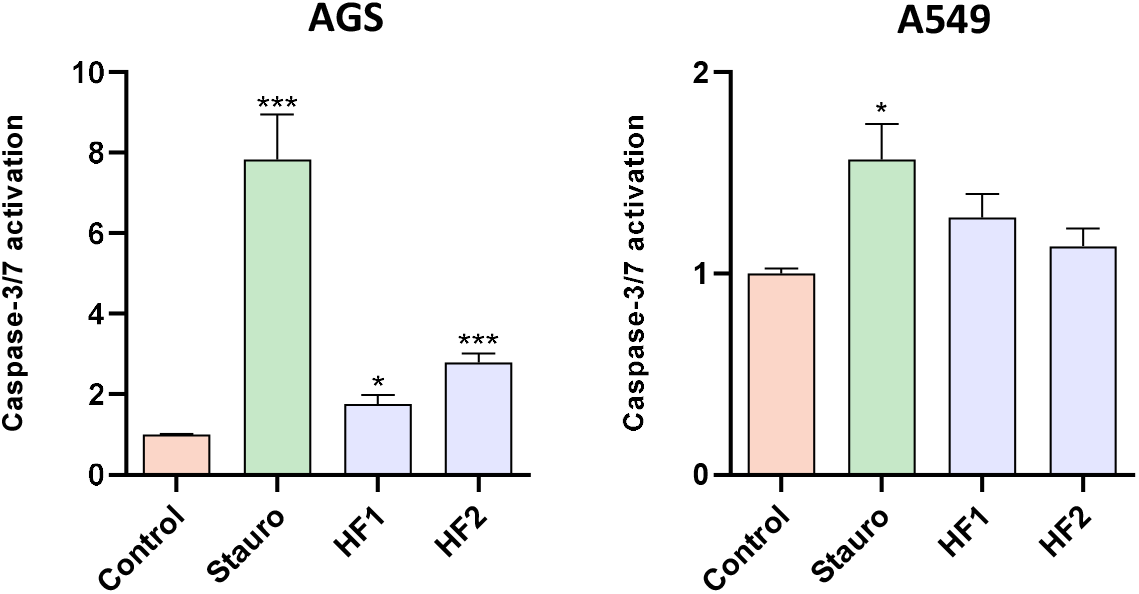
Fold increase in caspase 3/7 activity in AGS and A549 cells exposed to 6-(3-chloropropoxy)-flavanone (**HF1**, 100 μM) and 6-propoxyflavanone (**HF2**, 100 μM) for 24 h. Stauro: staurosporine, 100 nM). All results were normalized for DNA content. * *p < 0.05; ***p < 0.001.*

As for the caspase-3 activation assay results for the A549 cell line, the increase detected did not achieve statistical significance. In fact, even staurosporine elicited an effect less pronounced than the one detect for AGS cells (1.6 fold vs 8 fold, respectively), which confirms published data regarding the resistance of those cells. Taken together, the fact that marked reduction was found in the viability assay while no caspase-3 activation was detected could suggest that both 6-(3-chloropropoxy)-flavanone and 6-propoxyflavanone may exert a cytostatic effect in A549 cells instead of a cytotoxic one, in harmony with what had been found in morphological analysis of treated cells.

#### 3.4.6-(3-Chloropropoxy)-flavanone and 6-propoxyflavanone are selectively toxic towards cancer cells

At this point we had shown that 6-(3-chloropropoxy)-flavanone and 6-propoxyflavanone exhibit cytotoxicity towards both AGS and A549 cancer cells, and that this effect was necrosis-independent and involved caspase-3 activation. However, another desirable trait for cytotoxic drugs is their selectivity towards cancer cells, thus resulting in safer and more selective molecules that reduces their toxic effects towards other cell populations and, consequently, fewer side effects. In order to assess selectivity, we evaluated the effect of 6-(3-chloropropoxy)-flavanone and 6-propoxyflavanone using non-cancer cells, specifically the lung fibroblast cell line MRC-5, a benchmark in numerous toxicological studies [27,28]. As shown in **Figure 7**, neither 6-(3-chloropropoxy)-flavanone or 6-propoxyflavanone induced a significant loss in viability on MRC-5 cells. Moreover, since A549 is also a lung cell line, these two form a pair of cancer/non-cancer cell lines, which is useful to study the selectivity of a particular molecule and its activity on a specific organ or tissue.

**Figure 7.**
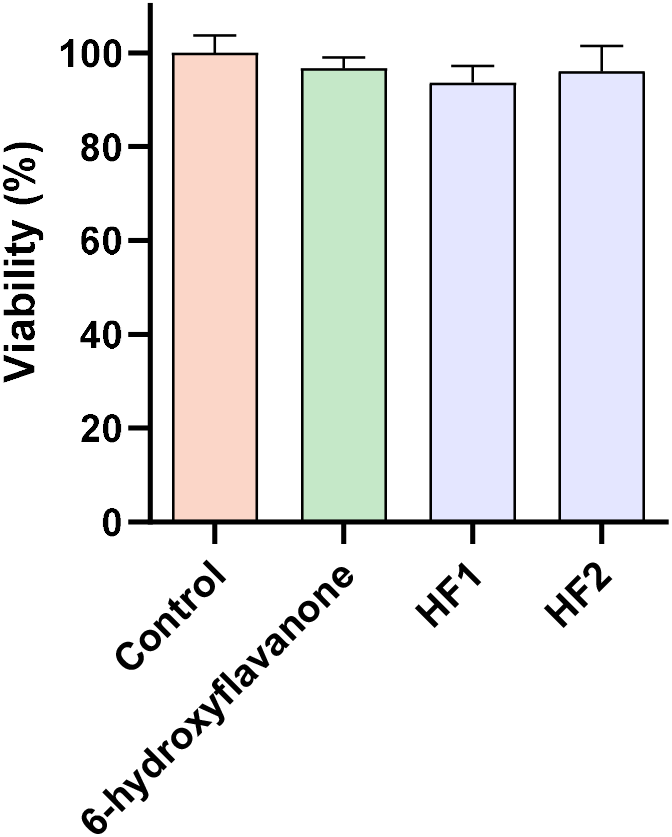
Viability of MRC-5 cells exposed to 6-hydroxyflavanone, 6-(3-chloropropoxy)-flavanone (**HF1**) and 6-propoxyflavanone (**HF2**) at 100 μM, after 24h of incubation.

6-Hydroxyflavanone has previously been shown to to exhibit no cytotoxic effects towards MRC-5 cells at 50 μM, as well as some of its derivatives, although these were oxime derivatives and not O-alkylated such as the ones presented herein [29]

## 4. Conclusion

In this work we explored the potential of structurally modified phenolics as an approach to the development of new anticancer drugs.

The cytotoxic activity of twenty three derivatives was tested on AGS and A549 cells, two widely used cell lines in pharmacological studies in the field of chemotherapy. The compounds were put through a screening of cytotoxic activity in order to gauge how many of these compounds were more potent than the parent molecules. Only two molecules were selected, namely **HF1** (6-(3-chloropropoxy)-flavanone) and **HF2** (6-propoxyflavanone). Both compounds exhibited a dose-dependent response, and did not cause any membrane damage at the concentrations tested.

Further investigation of the cell death processe taking place confirmed that these two derivatives of 6-hydroxyflavanone elicited cell death through apoptosis in AGS cells, namely activation of caspase-3. In the case of A549, a cytostatic effect is suggested. The selectivity profile assay conducted suggests these derivatives exhibit a selective anticancer activity, as they did not induce a significant loss of viability in non-cancer cells.

This work demonstrated that natural sources retain a diversified pool of potentially reactive chemical structures which can be modified to boost their medicinal applications.

## Credit authorship contribution statement

Conceptualization: David M. Pereira and Renato B. Pereira. Investigation: Pedro Olim, Maria José G. Fernandes, Carolina M. Natal, José R. A. Coelho; Data curation and formal analysis: David M. Pereira, Renato B. Pereira, M. Sameiro T. Gonçalves, Maria José G. Fernandes, Carolina M. Natal, José R. A. Coelho; Resources: David M. Pereira, M. Sameiro T. Gonçalves; Supervision: David M. Pereira, Renato B. Pereira, M. Sameiro T. Gonçalves, Maria José G. Fernandes, A. Gil Fortes; Writing – original draft: Pedro Olim; Writing – review & editing: David M. Pereira, Renato B. Pereira, M. Sameiro T. Gonçalves.

## Acknowledgements

The work was supported through projects UIDB/50006/2020, UIDP/50006/2020 and PTDC/ASP-AGR/30154/2017 (PO-CI-01-0145-FEDER-030154) of COMPETE 2020, funded by FCT (Fundação para a Ciência e Tecnologia, Portugal) /MCTES (Ministério da Ciência, Tecnologia e Ensino Superior) through national funds. Renato B. Pereira thanks FCT - Fundação para Ciência e a Tecnologia, I.P. for the funding through project PTDC-QUI/2870/2020. Fluorescence microscope acquired in the framework of project MILKQUA - H2020-PRIMA 2018—Section 2. The authors acknowledge also to FCT, and FEDER-COMPETE-QREN-EU for financial support to the research centre CQUM (UID/QUI/00686/2021).

The NMR spectrometer Bruker Avance III 400 is part of the National NMR Network and was purchased within the framework of the National Program for Scientific Re-equipment, contract REDE/1517/RMN/2005 with funds from POCI 2010 (FEDER) and FCT.

